# A genome-wide analysis of DNA methylation identifies a novel association signal for Lp(a) concentrations in the *LPA* promoter

**DOI:** 10.1101/627273

**Authors:** Stefan Coassin, Natascha Hermann-Kleiter, Margot Haun, Simone Wahl, Rory Wilson, Bernhard Paulweber, Sonja Kunze, Thomas Meitinger, Konstantin Strauch, Annette Peters, Melanie Waldenberger, Florian Kronenberg, Claudia Lamina

## Abstract

Lipoprotein(a) [Lp(a)] is a major cardiovascular risk factor, which is largely genetically determined by one major gene locus, the *LPA* gene. Many aspects of the transcriptional regulation of *LPA* are poorly understood and the role of epigenetics has not been addressed yet. Therefore, we conducted an epigenome-wide analysis of DNA methylation on Lp(a) levels in two population-based studies (total n=2208). We identified a CpG site in the *LPA* promoter which was significantly associated with Lp(a) concentrations. Surprisingly, the identified CpG site was found to overlap the SNP rs76735376. We genotyped this SNP de-novo in three studies (total n=7512). The minor allele of rs76735376 (1.1% minor allele frequency) was associated with increased Lp(a) values (p=1.01e-59) and explained 3.5% of the variation of Lp(a). Statistical mediation analysis showed that the effect on Lp(a) is rather originating from the base change itself and is not mediated by DNA methylation levels. This finding is supported by eQTL data from 153 liver tissue samples from the GTEx project, which shows a significant association of the rs76735376 minor allele with increased *LPA* expression. In summary, our data supports a functional role of rs76735376 in the regulation of *LPA* expression.

## Introduction

High Lipoprotein(a) [Lp(a)] concentrations represent a major risk factor for several cardiovascular diseases(1). Lp(a) concentrations are mostly genetically determined by one major gene locus, the *LPA* gene(2), which presents a peculiar structure consisting of several so-called kringle domains (kringle 4 type one to type 10 and kringle 5; abbreviated KIV-1 to -10 and KV). The KIV-2 domain (consisting of two exons) is encoded by a hypervariable copy number variation, which can be present in 1 to >40 copies per allele(1) and thus generates >40 isoforms of the protein [apolipoprotein(a)]. In Caucasians, low molecular weight isoforms (11-22 KIV repeats) are associated with 4-5 times higher Lp(a) than high molecular isoforms with >22 repeats.

The apo(a) isoform determined by the KIV-2 repeat number is the strongest contributor to Lp(a) concentration variance and explains ≈30-70 % of the total trait variance, with the complete *LPA* locus explaining up to 90%(2, 3), Interestingly, Lp(a) levels of the same-sized isoforms vary by 200-fold in the general population(1) but <2.5-fold within families(4), suggesting the existence of genetic variation strongly modifying the impact of the isoform. Indeed, genome-wide SNP-bound heritability has been recently estimated to be ≈49% in a German population(5).

Some high impact loss-of-function variants have been described(6–10) but regulatory variants have been elusive so far. The search for regulatory variants has so far focused on the promoter region(11–13) and a known enhancer region located 15 to 25 kb upstream of *LPA*(14), as well as on genome-wide association study efforts(5, 15, 16). Three variants in the enhancer region have been found to regulate *LPA* expression in reporter assays(14). Conversely, the promoter region of *LPA* is less well defined. Depending on whether a reference transcript with a leading non-coding first exon(11) (UCSC Genome browser hg38 annotation) or without it(12, 13) (NCBI NM_005577.2) was used, two different genomic regions located ≈4 kb apart from each other have been investigated as putative promoters(11). The most extensively studied region extends about 1.4 kb upstream of the translation start. Some regulatory variants(17, 18) and liver-specific regulatory elements(19) have been described in this region, as well as a pentanucleotide repeat (PNR) polymorphism with 5-12 repeats(12, 20). The latter is significantly associated with Lp(a) levels in Caucasians(21, 22). Early reports indicated a causal impact on promoter activity(12), which was not replicated in later studies(13). Since the association of the PNR with Lp(a) is, however, well replicated(21, 22), it is likely that the PNR is simply in linkage disequilibrium with a regulatory variant(22).

Epigenetic regulation presents an additional layer of genetic regulation acting via chemical modifications of either histones(23) or the DNA sequence itself(24). DNA methylation is a process by which methyl groups are added to the DNA molecule(24). 5-Methylcytosine can be activating, when occurring in gene bodies, or repressing, when occurring in promoters(24) (albeit the repressing action of promoter methylation extends into the 5’ region of the gene body(25)). DNA methylation may act by modifying the binding affinity of transcription factors(26) or recruiting silencing factors causing heterochromatinization(24). Variation in the methylation level can thus affect both binding affinity of transcription factors and chromatin organization.

Allele-specific methylation (ASM) at SNPs represents a special case of gene regulation via methylation. Up to 50,000, mostly cis-acting, ASM-SNPs have been reported(25, 27) so far. The large majority was located in CpG dinucleotides(28), where the SNP either creates or abolishes a CpG site (CpG-SNP). Such ASM-SNPs have been reported for e.g. inflammatory bowel disease(29), osteoarthritis susceptibility(30), atopic dermatitis(31), triglyceride levels(32) and diabetes(33). Even methylation close to a SNP site may already modulate the allele specific binding of transcription factors to the SNP site itself(34).

The impact of methylation on regulation of lipid phenotypes has been studied to a lesser extent than germline SNPs on these phenotypes. The studies performed so far reported clear associations with lipid phenotypes such as VLDL-C, HDL-C, LDL-C, triglycerides and total cholesterol and even lipid-related diseases (reviewed in(35)), but none of these studies has addressed Lp(a). Therefore we report here the first epigenome-wide methylation association study on Lp(a) concentrations.

## Material & Methods

### Study design and populations

The overall study design is outlined in Supplementary Figure 1. Three population-based studies were available for this analysis: The KORA F3, KORA F4 and SAPHIR studies. The KORA study (Cooperative health research in the region of Augsburg) consists of independent, non-overlapping, population-based samples from the general population living in the region of Augsburg, Southern Germany, and was conducted in the years 2004/2005 (KORA F3) and 2006 to 2008 (KORA F4)(36). A total of 3080 subjects with ages ranging from 32 to 81 years participated in the KORA F4 examination (stratified by sex-and age-groups). The genome-wide DNA methylation patterns were determined in a random subgroup of 1802 subjects (n=1724 in analysis dataset with non-missing values in all relevant variables). In KORA F3 (n=3184, age-range between 35 and 84 years), DNA methylation data were available from 484 subjects (comprised of 250 smokers and 250 non-smokers; 16 samples are missing due to data preprocessing or missing values in one of the relevant variables). For additional de novo genotyping, samples and data from 3080 KORA F3 and 2986 KORA F4 study participants were available, as well as samples from the SAPHIR study. For the SAPHIR study (Salzburg Atherosclerosis Prevention Program in subjects at High Individual Risk), participants were recruited by health screening programs in large companies in and around the city of Salzburg, Austria (n=1770, age-range 39-67 years). Genotypes and relevant phenotype data were available for 1446 participants. All study participants provided a signed informed consent and the studies were approved by their respective local ethics committee (Bayerische Landesärztekammer for the KORA studies, Ethical Committee of Salzburg for SAPHIR). The PNR was measured in a subset of the KORA F4 (n=2956) and SAPHIR (n=1428) studies.

### Genome-wide DNA methylation

Genome-wide DNA methylation patterns in the discovery and replication cohorts were analyzed using the Infinium HumanMethylation450 BeadChip Array (Illumina) and DNA derived from peripheral blood. KORA F3 and F4 samples were measured and processed separately (37). KORA F3/F4 samples were processed on 20 respectively 7 96-well plates in 9 respectively 4 batches; plate and batch effects were investigated using principal component analysis and eigenR2 analysis(38). DNA methylation data were preprocessed following the CPACOR pipeline(39). Background correction was performed using the R package minfi, v1.6.0(40). Signals with detection p-values (the probability of a signal being detected above the background) ≥ 0.01 were removed, as they indicate unreliable signals, as were signals summarized from fewer than three functional beads on the chip. Observations with less than 95% of CpG sites providing reliable signals were excluded.

To reduce the non-biological variability between observations, data were normalized using quantile normalization (QN) on the raw signal intensities: QN was performed on a stratification of the probe categories into 6 types, based on probe type and color channel, using the R package limma, v3.16.5(41).

The percentage of methylation at a given cytosine is reported as a beta-value, which is a continuous variable between 0 and 1 corresponding to the ratio of the methylated signal over the sum of the methylated and unmethylated signals of the particular cytosine site. Since it was the primary idea to detect the effect of differential methylation on Lp(a) and since probe binding might be affected by SNPs in the binding area, sites representing or being located in a 50 bp proximity to SNPs with a minor allele frequency (MAF) of at least 5% were excluded from the data set(42). Nine CpG sites in the *LPA* gene locus were present on the microarray (Supplementary Table 1).

### SNP validation and *LPA* pentanucleotide repeat genotyping

KORA F4 samples were genotyped using the Affymetrix Axiom chip array. Genotypes were called with the Affymetrix software and were annotated to NCBI build 37. Imputation was performed with IMPUTE v2.3.0 using the 1000G phase1 (v3) reference panel(43). Additional de novo genotyping for rs76735376 was performed in KORA F3, KORA F4 and the SAPHIR using a commercial Taqman assay (ThermoFisher Scientific).

Due to the low MAF, samples being heterozygous and homozygous for the rare allele were confirmed by Sanger sequencing using the primers 5’-TACAGGACAGAGACTAACT-3’ and 5’-GCATAGTATCAATCTTTCCG-3’. The PCR conditions were: Qiagen Taq DNA Polymerase (#201207), 0.2 mM dNTP, 0.5 μM primer, 1x Qiagen Q-Solution reagent, 40 ng input DNA amount. Cycling conditions were 94°C 3’ initial denaturation, 40 cycles of 94°C 30 sec, 60°C 30 sec, 72°C 30 sec, 72 °C 10 minutes final elongation. Sequencing was done using ABI BigDye 1.1 chemistry (Thermo Fisher Scientific Scientific, Waltham, MA, USA).

The *LPA* pentanucleotide repeat (PNR) is a microsatellite with 5-12 repeats of a TTTTA element located at hg19:chr6:161,086,617-161,086,663 (according to the RepeatMasker tool in the UCSC Genome Browser(44)) and thus only ≈50 bp downstream of rs76735376. It has been genotyped by fragment analysis in SAPHIR and KORA F4. Details of the PNR genotyping via fragment analysis are given in reference(20). In brief, a 181 bp PCR fragment encompassing the *LPA* pentanucleotide repeat was amplified in KORA F4 (data available in n=2956) and SAPHIR (data available in n=1429) in 384 well plates in a total volume of 5 μl with 2.5 μl Qiagen Multiplex PCR Plus Kit (Qiagen, Hilden, Germany) and 1 μl Qiagen Q-solution using a Yakima Yellow-labelled primer. Fragment length was detected by fluorescent capillary fragment analysis on an ABI 3730s Genetic Analyzer and all data analyzed using the GeneMapper software (all Thermo Fisher Scientific). Data was quality controlled for null allele overrepresentation and deviation from Hardy-Weinberg Equilibrium using Micro-Checker(45) (http://www.microchecker.hull.ac.uk/) and ARLEQUIN (http://cmpg.unibe.ch/software/arlequin3/)(46).

### Bisulfite sequencing

Bisulfite Sanger sequencing was used to validate the methylation signal at rs76735376 and map the methylation status of other CpG sites nearby in 8 selected samples from SAPHIR (4 homozygotes for the major allele and 4 heterozygotes). Supplementary Figure 2 provides an overview of the region with location of the PNR, the primers, the amplicons and the CpG affected by rs76735376, as well as the neighboring CpG sites covered (respectively not covered) by our bisulfite sequencing.

500 ng sample DNA were bisulfite treated using the kit EZ DNA Methylation-Lightning Kit (Zymo Research, Irvine, CA, USA) and eluted in two times 10 μl. 1 μl eluted converted DNA was used as template for PCR. The region was amplified using the DNA Polymerase of the Agena PCR Accessory and Enzyme Set (Agena Bioscience, San Diego, CA, USA). PCR conditions were: 0.4 u Agena DNA Polymerase, 0.5 mM dNTP, 0.2 μM primer, 2 mM MgCl_2_, 1 μl converted DNA. Cycling conditions were 94°C 4 minutes initial denaturation, 45 cycles of 94°C 20 sec, Annealing 30 sec, 72°C 60 sec, final elongation at 72 °C 3 minutes. Primers for bisulfite-converted DNA were designed using the Agena EpiTyper Software Suite (see Supplementary Table 2 for sequences and annealing temperature). All sequencing was done using ABI BigDye 1.1 chemistry.

### Measurement of Lp(a) concentrations and apo(a) isoform size

Lp(a) concentration were determined in plasma using the same standardized sandwich ELISA assay in all samples. The assay has been described in detail previously(10, 47). In brief, ELISA plates were coated with an affinity-purified rabbit anti-human apo(a) antibody and incubated with human plasma for 1 hour. After several washing steps, the bound Lp(a) was detected using a horseradish-peroxidase-conjugated monoclonal antibody (1A2; in 0.1% w/v casein, 1x PBS, pH 7.3) and the Blue Star TMB substrate (Adaltis, Guidonia Montecelio, IT) (30 minutes incubation at room temperature).

Lp(a) isoform was determined by Western blotting in all samples as described(10). In brief, 150 ng Lp(a) as determined in the ELISA were loaded on a 1.46*%* agarose gel with 0.08*%* SDS and separated for 18 h at 0.04 A (constant current). A size standard consisting of a mixture of five plasma samples collected from individuals expressing each only one apo(a) isoform (13, 19, 23, 27, 35 KIV repeats determined by Pulsed Field Gel Electrophoresis) was applied in every seventh well of the gel. The gel was subsequently blotted to a PVDF membrane (Immobilion-P, Millipore, Darmstadt, DE) by semi-dry blotting (Perfect Blue Semi Dry Blotter, VWR, Vienna, A). After blocking (85 mM NaCl, 10 mM TRIS, 0.2*%* Triton X-100, 1*%* BSA; ≥30 min at 37°C), the membrane was incubated with horseradish-peroxidase-conjugated 1A2 antibody (1A2 concentration was 168 ng/ml in Buffer C) for 2 h at room temperature on a shaker. After washing with TTBS (3x 15 min), ECL substrate (WesternBright Chemilumineszenz Spray, Biozym, Hessisch Oldendorf, DE) was added and signals recorded on an Amersham Hyperfilm™ ECL™ (GE Healthcare, Vienna, A).

In heterozygous individuals, the smaller (and thus the commonly Lp(a) concentration-determining) of the two alleles was used for isoform-adjustment of statistical models.

### Statistical methods

#### Genome wide methylation analysis

Linear regression models were applied to evaluate the association between DNA methylation beta values and Lp(a). Lp(a) levels were log-transformed to account for the skewed distribution. Age, sex and the first 20 principal components of the Illumina control probes (to adjust for technical confounding) were included as potential confounders. In addition, since whole blood DNA samples were used, cell heterogeneity had to be considered as a confounder. As no measured cell count information was available, sample-specific estimates of the proportion of the major white blood cell types were obtained using a statistical method described by Houseman et al(48). To correct for multiple comparisons, a Bonferroni corrected significance level of 1.03e-07 (=0.05/485512) was used. Genome-wide significant findings were replicated in KORA F3 using the same adjustment model as in the discovery stage. Fixed-effects inverse-variance meta-analysis was applied to summarize the findings in KORA F3 and F4. The analyses were repeated additionally adjusting for apo(a) isoforms or in subgroups of apo(a) isoforms.

#### SNP association and mediation analysis

Association of the identified SNP with log-transformed Lp(a) was evaluated in all three studies individually (KORA F3, KORA F4, SAPHIR) using a linear regression model and in all three studies combined using a linear mixed effects model with “study” as a random effect and adjusting for age and sex. The genotypes of the SNP were additively coded, that is, effect sizes refer to one copy of the minor allele. This analysis was repeated additionally adjusting for apo(a) isoforms or in subgroups of apo(a) isoforms. In addition to the models with log-transformed Lp(a) values as the dependent variable, which were used to ensure the normal-distribution assumption to derive the test-statistic and p-values, Lp(a) on its original scale was used. From these models, beta estimates and standard errors (se) were derived to ease interpretability of the effect sizes.

Due to the close vicinity of the cg-site/SNP with the well-known PNR polymorphism, the association of the PNR with Lp(a) concentrations was evaluated and whether this effect can be explained by the SNP or vice versa.

To assess whether the effect of the identified SNP on Lp(a) is mediated through methylation, a formal mediation analysis was applied using the R library “mediation”.

### Follow-up in GTEx and transcription factor binding site prediction

The effect of rs76735376 on *LPA* gene expression in native liver tissue was assessed using the GTEx(49) data (www.gtexportal.com), which contains expression quantitative trait locus (eQTL) data with whole genome SNP data and whole transcriptome RNA-Seq data for 153 liver samples. The correlation of rs76735376 genotype with *LPA* expression in liver was extracted from the GTEx dataset “Single Tissue-Specific All SNP Gene Associations” (available at https://storage.googleapis.com/gtex_analysis_v7/single_tissue_eqtl_data/all_snp_gene_associations/Liver.allpairs.txt.gz, accessed via https://gtexportal.org/home/datasets on August, 29^th^ 2018) using the 1000 Genomes SNP-ID “6_161086716” (chromosome and position of rs76735376 in hg19) and the Ensembl gene identifier for *LPA* “ENSG00000198670”.

TRANSFAC (http://genexplain.com/transfac/)(50), a database predicting the genomic binding sites and DNA-binding profiles of eukaryotic transcription factors, was used in order to determine transcription factor binding sites at the genomic region of rs76735376.

### Gel Mobility Shift Assay

To perform a pilot validation of the mediation analysis results, single-stranded oligonucleotides spanning the SNP site were synthesized by Eurofins MWG Operon and then annealed to generate double-stranded probes. The single-stranded oligonucleotide sequences for the C, T and 5-mC allele of rs76735376 respectively are shown in Supplementary Figure 3. 5-mC bases were introduced directly by the oligonucleotide provider during oligonucleotide synthesis. Annealing was performed by heating a mix of both oligos (2.5 μg in 100 μl with 1X annealing buffer: 20 mM TRIS ph 8.0, 1 mM EDTA, 50 mM NaCl) to 65° C for 5 minutes and slowly cooling down in ice water. Binding reactions for super shift assays were performed for 30 min at 4-8°C using the Binding Buffer B-1 (Active Motif 37480) together with Stabilizing Solution D (Active Motif 37488) containing either 5 μg human liver or embryonic kidney nuclear extracts (Active Motif 36042; 36033), 2 μg of either anti c/EBPbeta (*CEBPB*) (Santa Cruz Biotechnology; sc-150x), anti-Oct-1 (*POU2F1*) (Santa Cruz Biotechnology; sc232) plus anti-Oct-3/4 (*POU5F1*) (Santa Cruz Biotechnology; sc5279) or anti-Nfatc1 (*NFATC1*) as negative control (Santa Cruz Biotechnology; sc7294). Binding reactions with 3 × 10^5^ cpm of labeled probe was performed for 20 min at 4°-8°C using Binding Buffer C-1 (Active Motif 37484) together with Stabilizing Solution D (Active Motif 37488). Samples were separated on a 4% native polyacrylamide gel in 0.5 × TBE for 3 h at 250 V. For competition assays, 10-fold unlabeled cold oligonucleotides identical to the radioactively labeled non-methylated Oligo2 LPA_EMAS2 or the radioactively labeled methylated Oligo6 LPA_EMAS2 probes were added to the binding reaction in addition.

## Results

### Methylation analysis

Descriptive statistics of the participating studies are given in Supplementary Table 3. One CpG site (cg17028067) located in the promoter region of the *LPA* gene was significantly associated with Lp(a) in the discovery sample KORA F4 (p=6.04e-11, Supplementary Table 1). This site was replicated in KORA F3 with direction-consistent effect size (p=0.0001 in KORA F3, meta-analysis results: p=2.61e-14; Table 1).

**Table 1:**
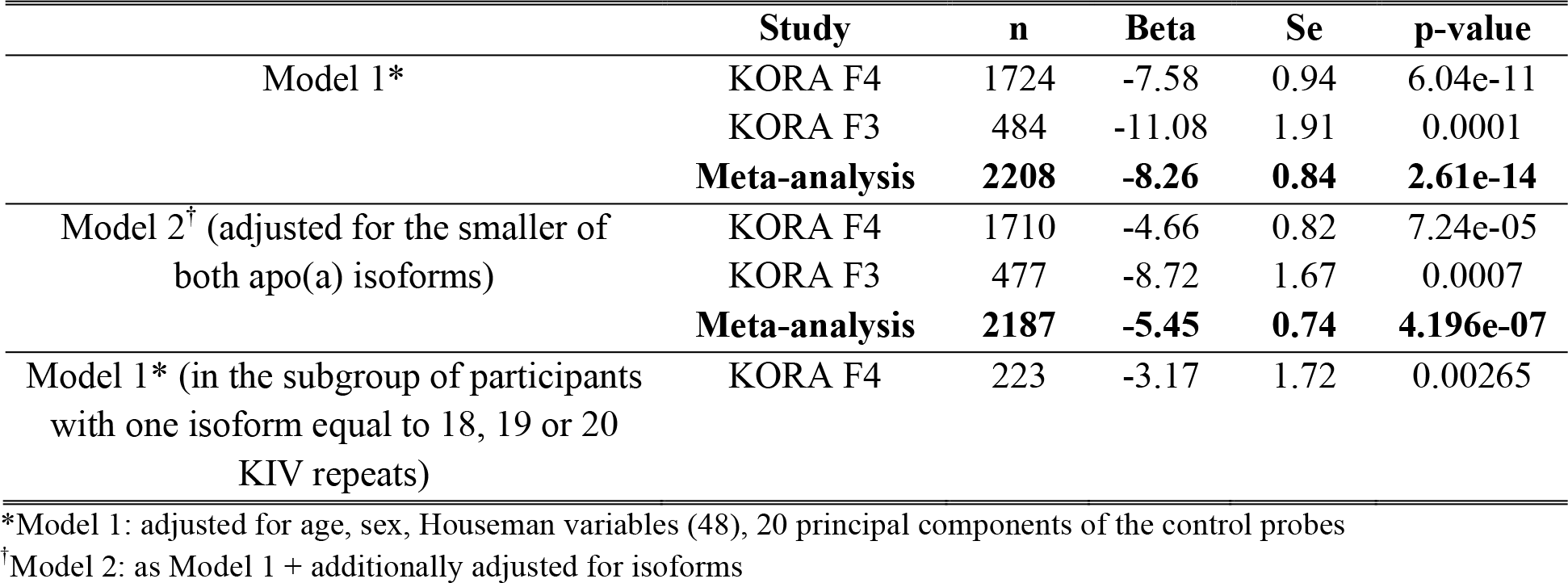
Results of the linear model of cg site cg17028067 on Lp(a). The p-value is derived from the log-transformed model, whereas beta estimate and standard error (se) are derived from the model on the original scale of Lp(a) to ease interpretability and refer to a change of 0.1 in methylation beta-value.

|  | Study | n | Beta | Se | p-value |
| --- | --- | --- | --- | --- | --- |
| Model 1* | KORA F4 | 1724 | -7.58 | 0.94 | 6.04e-11 |
|  | KORA F3 | 484 | -11.08 | 1.91 | 0.0001 |
|  | **Meta-analysis** | **2208** | **-8.26** | **0.84** | **2.61e-14** |
| Model 2<sup>†</sup> (adjusted for the smaller of both apo(a) isoforms) | KORA F4 | 1710 | -4.66 | 0.82 | 7.24e-05 |
|  | KORA F3 | 477 | -8.72 | 1.67 | 0.0007 |
|  | **Meta-analysis** | **2187** | **-5.45** | **0.74** | **4.196e-07** |
| Model 1* (in the subgroup of participants with one isoform equal to 18, 19 or 20 KIV repeats) | KORA F4 | 223 | -3.17 | 1.72 | 0.00265 |

\*Model 1: adjusted for age, sex, Houseman variables (48), 20 principal components of the control probes
<sup>†</sup>Model 2: as Model 1 + additionally adjusted for isoforms

Since low methylation beta-values of cg17028067 (mean ± sd in KORA F4 = 0.811 ± 0.068; in KORA F3: 0.782 ± 0.061) were occurring predominantly in low molecular weight apo(a) isoforms with 18 to 20 KIV repeats (Supplementary Figure 4), the analysis was enhanced as follows (Table 1): first additionally adjusting for the lower of both apo(a) isoforms and secondly analyzing only the subgroup of participants with one isoform equal to 18-20 repeats (only in KORA F4 due to low sample size in KORA F3). Although the association was partially attenuated after adjustment for apo(a) isoforms (p=4.20e-7, n=2187) and in the subgroup of isoforms 18-20 (p=0.00265, n=223), it still remained highly significant, indicating that the association is not only a consequence of a correlation of the SNP with small apo(a) isoforms.

### SNP genotyping

Surprisingly, the identified cg site was found to overlap a SNP (rs76735376). This C>T (reverse strand, i.e. transcribed strand of *LPA*) polymorphism is located exactly at the cytosine base of the CpG site cg17028067. The minor allele thus abolishes one potential methylation site by replacing the cytosine base of the CpG by a thymidine, thus effectively reducing the amount of available methylation target sites (Figure 1). This SNP had not been genotyped directly with the earlier SNP microarray but had been imputed with a low imputation quality (imputation quality info =0.4) and a reported very low minor allele frequency of ≈0.1%. Therefore, this cg site had not been removed from the primary analysis dataset when SNP positions were masked. Even more surprising, subsequent de novo genotyping and confirmatory Sanger sequencing in KORA F3, KORA F4 and SAPHIR revealed that this SNP was indeed considerably more frequent than reported in the imputed data with a true MAF of 1.1% and a carrier frequency of 2.2% (Supplementary Table 4).

**Figure 1:**
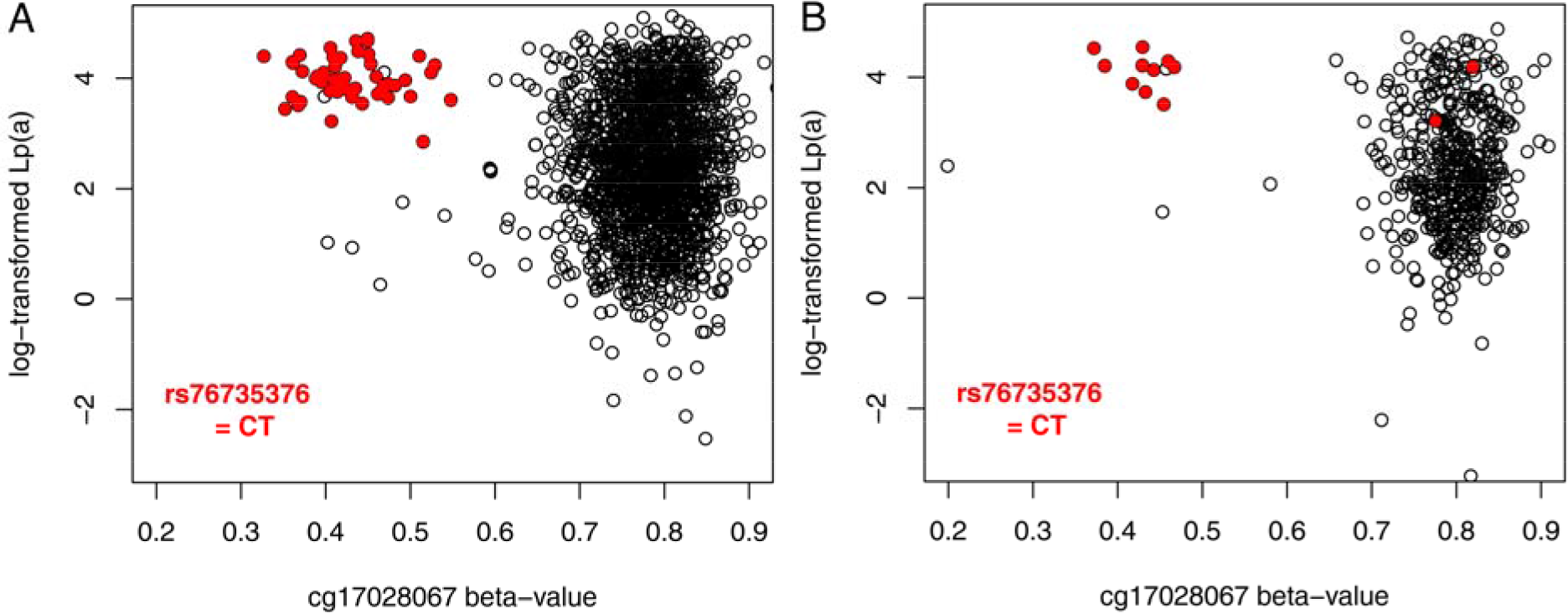
Relationship of methylation level at cg17028067 (x axis), Lp(a) values (log-transformed (y axis) and rs76735376 genotype in the KORA F4 (panel A) and KORA F3 study (panel B). heterozygous carriers of rs76735376 show roughly halved methylation levels and higher Lp(a) levels with less variance.

### Validation by genotyping and bisulfite sequencing

The methylation of the region was validated by bisulfite sequencing in 8 samples of the SAPHIR study (4 homozygotes for the major allele of rs76735376 and 4 heterozygotes, Supplementary Figure 5A). Besides cg17028067 nine additional CpG sites nearby cg17028067 were all nearly 100% methylated (Supplementary Figure 2). None of these sites were represented on the HumanMethylation450 BeadChip array. The methylation level of the cg sites nearby was independent from the status of cg17028067. This indicates that the whole region might contain CpG dinucleotides, which are target of DNA methylation and can thus potentially affect *LPA* expression. In heterozygous carriers of the SNP rs76735376 the methylation at cg17028067 was reduced by 50% as would be expected (Supplementary Figure 5B).

### SNP association analysis

The minor allele of rs76735376 was associated with increased Lp(a) values (Figure 2) with consistent effect sizes in all three cohorts (p=1.01e-59, Table 2). Each copy of the minor allele was associated with an increase of Lp(a) concentrations by 37 mg/dL. The SNP explained 3.45% of the variance of Lp(a) concentrations. One part of the SNP effect seems to be due to correlation with apo(a) isoforms. As already observed for the methylation levels, isoforms 18-20 are overrepresented in carriers of the minor allele (T) (Supplementary Figure 6). The effect size is reduced in a model adjusted for isoforms and one third to one quarter as high in subgroups with one apo(a) isoform equal to 18, 19 or 20 repeats. However, rs76735376 still explains about 3.2% of the variance of Lp(a) in the latter subgroup (p=9.71e-08, Table 2 and Figure 2).

**Figure 2:**
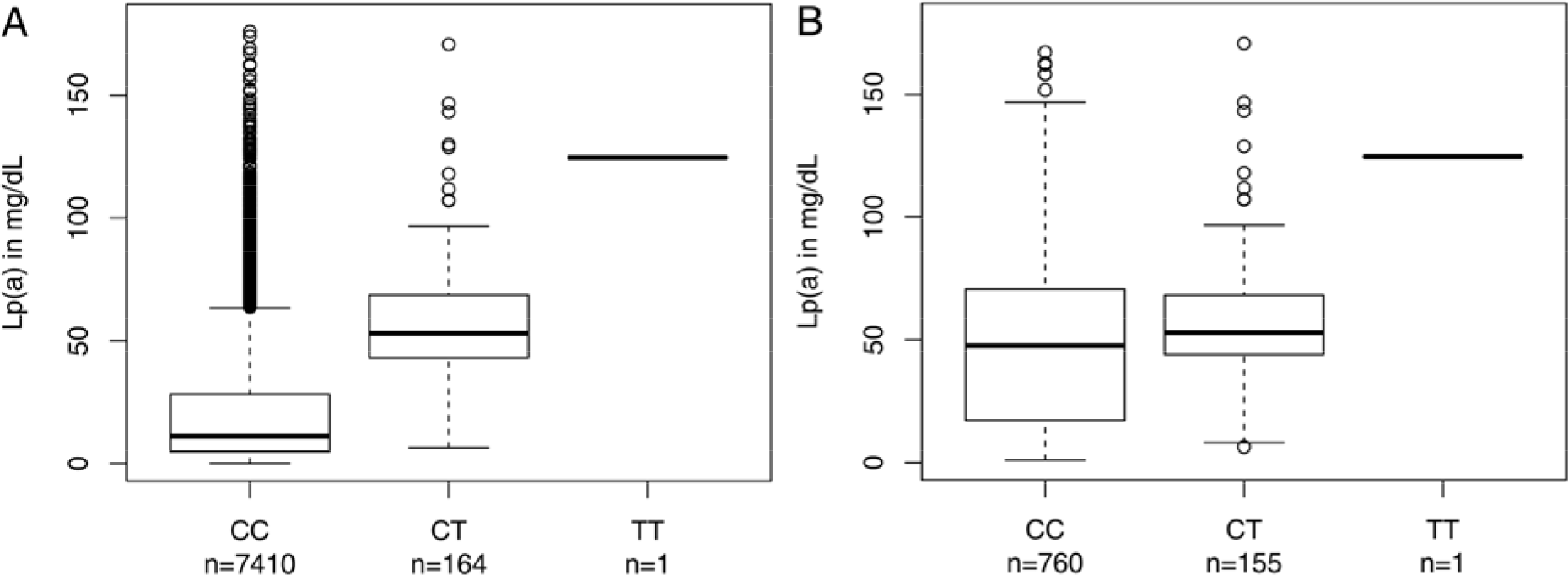
Distribution of Lp(a) stratified for genotypes of SNP rs76735376 in all participants of all three cohorts (panel A) and in subgroup of participants with one apo(a) isoform equal to 18, 19 or 20 repeats (panel B).

**Table 2:** Association analysis of rs76735376 on Lp(a) concentrations adjusted for age and sex. The p-value and variance explained are derived from the log-transformed model, whereas beta estimate and standard error (se) are derived from the model on the original scale of Lp(a) to ease interpretability and refer to the minor allele (T) of rs76735376.

| Study | Results of linear regression |  |  |  |
| --- | --- | --- | --- | --- |
|  | beta | se | p-value | Variance explained |
| **In all participants:** |  |  |  |  |
| KORA F4 | 36.42 | 2.81 | 3.23e-25 | 0.0384 |
| KORA F3 | 38.06 | 3.14 | 2.89e-27 | 0.0342 |
| SAPHIR | 36.84 | 5.23 | 9.81e-11 | 0.0273 |
| **Combined\*** | **37.12** | **1.96** | **1.01e-59** | **0.0346<sup>†</sup>** |
| **Additionally adjusted for the small isoform:** |  |  |  |  |
| KORA F4 | 19.65 | 2.49 | 1.37e-10 | 0.0062 |
| KORA F3 | 22.23 | 2.77 | 4.70e-09 | 0.0109 |
| SAPHIR | 20.11 | 4.62 | 8.09e-05 | 0.0104 |
| **Combined\*** | **20.71** | **1.73** | **1.73e-21** | **0.0089<sup>†</sup>** |
| **In participants with one isoform 18/19/20:** |  |  |  |  |
| KORA F4 | 6.42 | 4.16 | 0.0049 | 0.0440 |
| KORA F3 | 14.13 | 4.60 | 5.87e-05 | 0.0217 |
| SAPHIR | 8.61 | 7.49 | 0.0151 | 0.0289 |
| **Combined\*** | **9.44** | **2.86** | **9.71e-08** | **0.0322<sup>†</sup>** |

\*Linear mixed effects model with “study” as random effect; <sup>†</sup>weighted average of study-specific variance explained

### Association with the pentanucleotide-repeat (PNR) polymorphism

Since rs76735376 is in close vicinity to the well-known *LPA* pentanucleotide-repeat (PNR) polymorphism(12, 20–22), we evaluated in KORA F4 and SAPHIR, whether the association of the PNR with Lp(a) concentrations can be explained by rs76735376 or vice versa. In both studies, 89% of all participants present one allele with 8 repeats of this PNR, whereas all carriers of the minor allele of rs76735376 have this repeat number at least once (Supplementary Table 5, p=0.01, Fisher’s exact test in both studies combined). However, in a mixed effects model using data from both studies, the effect of the PNR on log-lp(a) (beta= −0.15, se=0.04, p=0.0003) is mediated only marginally by rs76735376 (results of PNR adjusted for rs76735376: beta= −0.14, se=0.04, p=0.0012). The SNP-effect is not affected at all by adding the PNR to the model (effect of SNP only in both studies on log-lp(a): beta= 1.55, se=0.12, p=3.82e-37; Adjusted for PNR: beta= 1.53, se=0.12, p=1.62e-36).

### Mediation analysis

Since rs76735376 was identified actually by chance in a genome-wide methylation analysis, we investigated whether the effect of the SNP on Lp(a) is mediated through the methylation signal. A formal mediation analysis revealed no significant causal mediation effect of the methylation level of cg17028067 on log-Lp(a) (p=0.13, Supplementary Figure 7). The total effect of the SNP on Lp(a) is only marginally altered by inclusion of the beta-value of cg17028067 in the model. In a regression model with both, rs76735376 and cg17028067, only the SNP remains significant. This might indicate that the true causal factor is the base change at rs76735376 rather than the methylation level at cg17028067.

### Effect of rs76735376 on *LPA* transcription

*LPA* is expressed exclusively in liver tissue, which is hard to obtain for a large number of samples, especially if they need to be controlled for defined rare *LPA* genotypes. To assess whether rs76735376 indeed affects *LPA* expression in vivo, we therefore performed a lookup in the data of the GTEx consortium(51). GTEx is a multinational project providing both genome-wide SNP data and complete transcriptome data for >10,000 individuals and >70 different tissues, including data from 153 liver samples. The minor allele rs76735376 was significantly associated with increased *LPA* expression in liver tissue (beta(se)=+0.44 (0.20), p=0.0369) despite only 11 heterozygous carriers in the data. Accordingly, bioinformatic transcription factor binding analysis using TRANSFAC(50) revealed a direct DNA transcription factor binding site for Oct factors *POU2F1* and *POU5F1* (also known as Oct1 and Oct3/4) within the T -allele of the rs76735376 SNP but not the C- or 5-mC allele (Supplementary Figure 3). Moreover, 5 bases besides the Oct binding site, a *CEBPB* was predicted (Supplementary Figure 3). Pilot experiments using electrophoretic mobility shift assays (EMSA) with probes encompassing the SNP site with C, 5-mC and T allele oligonucleotides suggests that *CEBPB* but not the Oct factors *POU2F1* and *POU5F1* indeed bind to both the C and 5-mC allele, but not to the T-allele (Supplementary Figure 8).

## Discussion

Lp(a) is a major genetically determined cardiovascular risk factor. Research on Lp(a) has experienced two revivals: the first by genetic studies providing strong support for a causal association with cardiovascular outcomes and the second, more recently, due to Lp(a)-lowering agents becoming available in the form of PCSK9 inhibitors and mRNA antisense therapy. Despite the strong impact of Lp(a) on CVD, many controversies and uncertainties remain about its metabolism and regulation(52). The association of Lp(a) concentrations with the apo(a) isoform size is well accepted, but far from being simple and linear(1, 10). Even within the same isoform group, levels may vary by 200-fold(4), suggesting the existence of modifier variants which act on top of the apo(a) isoforms. Some of these have been identified(7, 53, 54), but they all act via disturbing protein synthesis.

Regulatory variants and elements are less defined. However, understanding the transcriptional regulation of *LPA* might be highly relevant for future drug development. We performed the first genome-wide DNA methylation study for Lp(a). This identified the SNP rs76735376 located in a CpG site in the *LPA* promoter. Each minor allele abolishes a methylation site in a CpG dinucleotide and is associated with an increase in Lp(a) of about 37 mg/dL and 9 mg/dL when restricted to individuals with 18-20 KIV repeats who are the main carriers of this methylation site. To dissect the effect of the base from the methylation site we performed a statistical mediation analysis which indicated that the effect observed in the association study is driven by the base change and not by the methylation status of the locus. This was supported by pilot EMSA experiments using the three possible allele states C, 5m-C and T), which showed no clearly defined difference between the C and the 5m-C allele, but a difference between the C and T alleles.

The location in the promoter suggests an effect on the transcriptional regulation of *LPA.* While Lp(a) is exclusively expressed in the liver and native tissue is hard to obtain in numbers sufficient to account for the low MAF of rs76735376. The recent large-scale multi-tissue transcriptome and eQTL profiling initiative GTEx, however, provides a convenient way to still investigate the effects of rs76735376 on *LPA* expression. Indeed, a look-up for in the GTEx raw data revealed that the rs76735376 T-allele is associated with significantly increased *LPA* expression, which is in line with the observed increase of Lp(a) concentrations in these individuals.

Interestingly, rs76735376 was located close to the *LPA* pentanucleotide repeat (PNR). This led us to speculate that it might be the causal factor underlying the previously reported(12), but not clearly replicated(13) impact of the PNR on *LPA* expression. This was, however, not the case as the association signal at rs76735376 was only marginally affected by including the PNR in the model and vice versa. Rs76735376 therefore represents a novel, independent SNP in the *LPA* promoter strongly associated with Lp(a) concentrations. Although it is rare, it explains more than 3% of the Lp(a) variance, which is remarkable high compared to other complex phenotypes.

Of note, all carriers of the minor allele of rs76735376 carried a PNR allele with 8 copies and virtually all minor allele carriers presented an apo(a) allele with 19 to 20 KIV repeat (Supplementary Figure 6). Rs76735376 therefore is in line with several other putative functional SNPs in *LPA*, which are confined to certain isoform ranges(18, 53, 55). This underscores the complex genetic make-up of the Lp(a) trait, consisting of an intricate interplay between the KIV-2 repeat number, transcriptional regulation by enhancer and promoter regions, modifier SNPs, which may be in turn confined only to certain isoform sizes, and trans-regulating factors like apoE(5, 56). Given the large impact of the underlying apo(a) isoform on Lp(a) levels, a full assessment of variants influencing Lp(a) is likely to require complete knowledge about the isoforms of the investigated samples. Failure to account for the isoforms has been shown to even potentially mask true genetic effects(18, 53).

Our study also highlights an important technical message for the community. Indeed, when we originally set out to investigate the impact of DNA methylation on Lp(a), the quality control procedure was supposed to purge any CpG sites affected by a SNP by using imputed 1000Genomes genotype data filtered at 5% MAF to focus only on differential methylation but not common ASM-SNPs. However, rs76735376 escaped filtering because it was badly imputed and thus incorrectly reported with a MAF of 0.1%. Since such a low MAF could not have triggered the association signal observed, we genotyped it de-novo, revealing a ten times higher true MAF. This MAF is now confirmed also by new reference datasets that became available only very recently (like TOPMed(57), https://bravo.sph.umich.edu/freeze5/hg38/).

Our study presents some limitations which need to be mentioned. Firstly, the methylation dataset was derived from peripheral blood, while *LPA* expression is limited to hepatic tissue. This is an unavoidable limitation of population-scale genome wide methylation analyses since hepatic tissue is not accessible at the sample numbers required for genome-wide studies. This applies even more to Lp(a), where a full depiction of the isoform variability is required for proper analysis and some functional variants are rare. Indeed, even the GTEx initiative, which contains data from ≈10,000 individuals and >70 tissues, reports only 175 liver samples (as of April 2019; thereof only 153 having also SNP genotype data available). Nevertheless, while methylation is clearly a tissue-specific marker, studies also have shown that a consistent portion of the DNA methylation patterns is still conserved across tissues(58, 59). Therefore, while the use of blood-derived DNA as surrogate for hepatic methylation may have led to missing some very strictly tissue-specific regulators, we can be confident that a consistent portion of methylation-based regulation has still been assayed. Secondly, our dataset was of Caucasian origin. Given the known differences in allele frequencies and effects of *LPA* SNPs between ethnicities, investigations in other populations are required to establish the meaning of our findings in other ethnicities(60, 61). Finally, functional and fine mapping studies that integrate tissue specific transcriptomics and epigenomics data with Lp(a) phenotype data are finally warranted to fully understand the contribution of this SNP to the Lp(a) trait.

In conclusion, this study represents a first step assessing the epigenetic regulation of Lp(a). We showed that the analyzed region of the Lp(a) promoter is subjected to DNA methylation and the identification of rs76735376 located at the CpG site at cg17028067 adds a novel variant to the catalog of genetic variants affecting Lp(a) concentrations. Extending our association results, we found an association between rs76735376 and *LPA* gene expression levels in public tissue data. Although the elucidation of the precise mechanisms and biological relevance of our observations will require dedicated in-depth studies, investigations on Lp(a) regulation might be fruitful and lead to novel therapeutic options.

## Abbreviations

*LPA*: lipoprotein(a) gene
Lp(a): lipoprotein(a) particle
EMSA: Electrophoretic mobility shift assays
PNR: Penta-Nucleotide-repeat

## Acknowledgements

**Sources of funding**

The study was supported by the Austrian Science Fund (FWF) Project Number P266600-B13 to CL. The KORA study was initiated and financed by the Helmholtz Zentrum München – German Research Center for Environmental Health, which is funded by the German Federal Ministry of Education and Research (BMBF) and by the State of Bavaria. This work was supported by a grant (WA 4081/1-1) from the German Research Foundation.

## References

1. Kronenberg, F., and G. Utermann. 2013. Lipoprotein(a): resurrected by genetics. J. Intern. Med. 273: 6–30.

2. Boerwinkle, E., C. C. Leffert, J. Lin, C. Lackner, G. Chiesa, and H. H. Hobbs. 1992. Apolipoprotein(a) gene accounts for greater than 90% of the variation in plasma lipoprotein(a) concentrations. J Clin Invest. 90: 52–60.

3. Schmidt, K., H. G. Kraft, W. Parson, and G. Utermann. 2006. Genetics of the Lp(a)/apo(a) system in an autochthonous Black African population from the Gabon. Eur. J. Hum. Genet. 14: 190–201.

4. Perombelon, Y. F. N., A. K. Soutar, and B. L. Knight. 1994. Variation in lipoprotein(a) concentration associated with different apolipoprotein(a) alleles. J. Clin. Invest. 93: 1481–1492.

5. Mack, S., S. Coassin, R. Rueedi, N. A. Yousri, I. Seppälä, C. Gieger, S. Schönherr, L. Forer, G. Erhart, P. Marques-Vidal, J. S. Ried, G. Waeber, S. Bergmann, D. Dähnhardt, A. Stöckl, O. T. Raitakari, M. Kähönen, A. Peters, T. Meitinger, K. Strauch, KORA-Study Group, L. Kedenko, B. Paulweber, T. Lehtimäki, S. C. Hunt, P. Vollenweider, C. Lamina, and F. Kronenberg. 2017. A genome-wide association meta-analysis on lipoprotein (a) concentrations adjusted for apolipoprotein (a) isoforms. J. Lipid Res. 58: 1834–1844.

6. Ogorelkova, M. 1999. Molecular basis of congenital lp(a) deficiency: a frequent apo(a) “null” mutation in caucasians. Hum. Mol. Genet. 8: 2087–2096.

7. Parson, W., H. G. Kraft, H. Niederstätter, A. W. Lingenhel, S. Köchl, F. Fresser, and G. Utermann. 2004. A common nonsense mutation in the repetitive Kringle IV-2 domain of human apolipoprotein(a) results in a truncated protein and low plasma Lp(a). Hum. Mutat. 24: 474–80.

8. Lim, E. T., P. Würtz, A. S. Havulinna, P. Palta, T. Tukiainen, K. Rehnström, T. Esko, R. Mägi, M. Inouye, T. Lappalainen, Y. Chan, R. M. Salem, M. Lek, J. Flannick, X. Sim, A. Manning, C. Ladenvall, S. Bumpstead, E. Hämäläinen, K. Aalto, M. Maksimow, M. Salmi, S. Blankenberg, D. Ardissino, S. Shah, B. Horne, R. McPherson, G. K. Hovingh, M. P. Reilly, H. Watkins, A. Goel, M. Farrall, D. Girelli, A. P. Reiner, N. O. Stitziel, S. Kathiresan, S. Gabriel, J. C. Barrett, T. Lehtimäki, M. Laakso, L. Groop, J. Kaprio, M. Perola, M. I. McCarthy, M. Boehnke, D. M. Altshuler, C. M. Lindgren, J. N. Hirschhorn, A. Metspalu, N. B. Freimer, et al. 2014. Distribution and medical impact of loss-of-function variants in the Finnish founder population. PLoS Genet. 10: e1004494.

9. Emdin, C. A., A. V. Khera, P. Natarajan, D. Klarin, H.-H. Won, G. M. Peloso, N. O. Stitziel, A. Nomura, S. M. Zekavat, A. G. Bick, N. Gupta, R. Asselta, S. Duga, P. A. Merlini, A. Correa, T. Kessler, J. G. Wilson, M. J. Bown, A. S. Hall, P. S. Braund, N. J. Samani, H. Schunkert, J. Marrugat, R. Elosua, R. McPherson, M. Farrall, H. Watkins, C. Willer, G. R. Abecasis, J. F. Felix, R. S. Vasan, E. Lander, D. J. Rader, J. Danesh, D. Ardissino, S. Gabriel, D. Saleheen, S. Kathiresan, CHARGE–Heart Failure Consortium, and CARDIoGRAM Exome Consortium. 2016. Phenotypic Characterization of Genetically Lowered Human Lipoprotein(a) Levels. J. Am. Coll. Cardiol. 68: 2761–2772.

10. Erhart, G., C. Lamina, T. Lehtimäki, P. Marques-Vidal, M. Kähönen, P. Vollenweider, O. T. Raitakari, G. Waeber, B. Thorand, K. Strauch, C. Gieger, T. Meitinger, A. Peters, F. Kronenberg, and S. Coassin. 2018. Genetic Factors Explain a Major Fraction of the 50% Lower Lipoprotein(a) Concentrations in Finns. Arterioscler. Thromb. Vasc. Biol. 38: 1230–1241.

11. Chennamsetty, I., T. Claudel, K. M. Kostner, A. Baghdasaryan, D. Kratky, S. Levak-Frank, S. Frank, F. J. Gonzalez, M. Trauner, and G. M. Kostner. 2011. Farnesoid X receptor represses hepatic human APOA gene expression. J. Clin. Invest. 121: 3724–34.

12. Wade, D. P., J. G. Clarke, G. E. Lindahl, A. C. Liu, B. R. Zysow, K. Meer, K. Schwartz, and R. M. Lawn. 1993. 5’ control regions of the apolipoprotein(a) gene and members of the related plasminogen gene family. Proc. Natl. Acad. Sci. U. S. A. 90: 1369–73.

13. Bopp, S., S. Köchl, F. Acquati, P. Magnaghi, A. Pethö-Schramm, H. G. Kraft, G. Utermann, H. J. Müller, and R. Taramelli. 1995. Ten allelic apolipoprotein[a] 5’ flanking fragments exhibit comparable promoter activities in HepG2 cells. J. Lipid Res. 36: 1721–8.

14. Puckey, L. H., and B. L. Knight. 2003. Sequence and functional changes in a putative enhancer region upstream of the apolipoprotein(a) gene. Atherosclerosis. 166: 119–27.

15. Trégouët, D.-A., I. R. König, J. Erdmann, A. Munteanu, P. S. Braund, A. S. Hall, A. Grosshennig, P. Linsel-Nitschke, C. Perret, M. DeSuremain, T. Meitinger, B. J. Wright, M. Preuss, A. J. Balmforth, S. G. Ball, C. Meisinger, C. Germain, A. Evans, D. Arveiler, G. Luc, J.-B. Ruidavets, C. Morrison, P. van der Harst, S. Schreiber, K. Neureuther, A. Schäfer, P. Bugert, N. E. El Mokhtari, J. Schrezenmeir, K. Stark, D. Rubin, H.-E. Wichmann, C. Hengstenberg, W. Ouwehand, A. Ziegler, L. Tiret, J. R. Thompson, F. Cambien, H. Schunkert, and N. J. Samani. 2009. Genome-wide haplotype association study identifies the SLC22A3-LPAL2-LPA gene cluster as a risk locus for coronary artery disease. Nat. Genet. 41: 283–5.

16. Li, J., L. A. Lange, J. Sabourin, Q. Duan, W. Valdar, M. S. Willis, Y. Li, J. G. Wilson, and E. M. Lange. 2015. Genome- and exome-wide association study of serum lipoprotein (a) in the Jackson Heart Study. J. Hum. Genet. 60: 755–61.

17. Zysow, B. R., G. E. Lindahl, D. P. Wade, B. L. Knight, and R. M. Lawn. 1995. C/T polymorphism in the 5’ untranslated region of the apolipoprotein(a) gene introduces an upstream ATG and reduces in vitro translation. Arter. Thromb Vasc Biol. 15: 58–64.

18. Kraft, H. G., M. Windegger, H. J. Menzel, and G. Utermann. 1998. Significant impact of the +93 C/T polymorphism in the apolipoprotein(a) gene on Lp(a) concentrations in Africans but not in Caucasians: confounding effect of linkage disequilibrium. Hum. Mol. Genet. 7: 257–64.

19. Wade, D. P., G. E. Lindahl, and R. M. Lawn. 1994. Apolipoprotein(a) gene transcription is regulated by liver-enriched trans-acting factor hepatocyte nuclear factor 1 alpha. J. Biol. Chem. 269: 19757–65.

20. Lamina, C., M. Haun, S. Coassin, A. Kloss-Brandstätter, C. Gieger, A. Peters, H. Grallert, K. Strauch, T. Meitinger, L. Kedenko, B. Paulweber, and F. Kronenberg. 2014. A systematic evaluation of short tandem repeats in lipid candidate genes: riding on the SNP-wave. PLoS One. 9: e102113.

21. Mooser, V., F. P. Mancini, S. Bopp, A. Pethö-Schramm, R. Guerra, E. Boerwinkle, H.-J. Müller, and H. H. Hobbs. 1995. Sequence polymorphisms in the apo(a) gene associated with specific levels of Lp(a) in plasma. Hum. Mol. Genet. 4: 173–181.

22. Trommsdorff, M., S. Köchl, a Lingenhel, F. Kronenberg, R. Delport, H. Vermaak, L. Lemming, I. C. Klausen, O. Faergeman, and G. Utermann. 1995. A pentanucleotide repeat polymorphism in the 5’ control region of the apolipoprotein(a) gene is associated with lipoprotein(a) plasma concentrations in Caucasians. J. Clin. Invest. 96: 150–7.

23. ENCODE Project Consortium. 2012. An integrated encyclopedia of DNA elements in the human genome. Nature. 489: 57–74.

24. Jones, P. A. 2012. Functions of DNA methylation: islands, start sites, gene bodies and beyond. Nat. Rev. Genet. 13: 484–492.

25. Schultz, M. D., Y. He, J. W. Whitaker, M. Hariharan, E. A. Mukamel, D. Leung, N. Rajagopal, J. R. Nery, M. A. Urich, H. Chen, S. Lin, Y. Lin, I. Jung, A. D. Schmitt, S. Selvaraj, B. Ren, T. J. Sejnowski, W. Wang, and J. R. Ecker. 2015. Human body epigenome maps reveal noncanonical DNA methylation variation. Nature. 523: 212–216.

26. Yin, Y., E. Morgunova, A. Jolma, E. Kaasinen, B. Sahu, S. Khund-Sayeed, P. K. Das, T. Kivioja, K. Dave, F. Zhong, K. R. Nitta, M. Taipale, A. Popov, P. A. Ginno, S. Domcke, J. Yan, D. Schübeler, C. Vinson, and J. Taipale. 2017. Impact of cytosine methylation on DNA binding specificities of human transcription factors. Science (80-.). 356.

27. Schalkwyk, L. C., E. L. Meaburn, R. Smith, E. L. Dempster, A. R. Jeffries, M. N. Davies, R. Plomin, and J. Mill. 2010. Allelic Skewing of DNA Methylation Is Widespread across the Genome. Am J Hum Genet. 86: 196–212.

28. Shoemaker, R., J. Deng, W. Wang, and K. Zhang. 2010. Allele-specific methylation is prevalent and is contributed by CpG-SNPs in the human genome. Genome Res. 20: 883–9.

29. John, G., J. P. Hegarty, W. Yu, A. Berg, D. M. Pastor, A. A. Kelly, Y. Wang, L. S. Poritz, S. Schreiber, W. A. Koltun, and Z. Lin. 2011. NKX2-3 variant rs11190140 is associated with IBD and alters binding of NFAT. Mol. Genet. Metab.

30. Reynard, L. N., C. Bui, C. M. Syddall, and J. Loughlin. 2014. CpG methylation regulates allelic expression of GDF5 by modulating binding of SP1 and SP3 repressor proteins to the osteoarthritis susceptibility SNP rs143383. Hum. Genet. 133: 1059–73.

31. Kumar, D., K. J. Puan, A. K. Andiappan, B. Lee, G. H. A. Westerlaken, D. Haase, R. Melchiotti, Z. Li, N. Yusof, J. Lum, G. Koh, S. Foo, J. Yeong, A. C. Alves, J. Pekkanen, L. D. Sun, A. Irwanto, B. P. Fairfax, V. Naranbhai, J. E. A. Common, M. Tang, C. K. Chuang, M.-R. Jarvelin, J. C. Knight, X. Zhang, F. T. Chew, S. Prabhakar, L. Jianjun, D. Y. Wang, F. Zolezzi, M. Poidinger, E. B. Lane, L. Meyaard, and O. Rötzschke. 2017. A functional SNP associated with atopic dermatitis controls cell type-specific methylation of the VSTM1 gene locus. Genome Med. 9: 18.

32. de Toro-Martín, J., F. Guénard, A. Tchernof, Y. Deshaies, L. Pérusse, S. Biron, O. Lescelleur, L. Biertho, S. Marceau, and M.-C. Vohl. 2016. A CpG-SNP Located within the ARPC3 Gene Promoter Is Associated with Hypertriglyceridemia in Severely Obese Patients. Ann. Nutr. Metab. 68: 203–12.

33. Dayeh, T. A., A. H. Olsson, P. Volkov, P. Almgren, T. Rönn, and C. Ling. 2013. Identification of CpG-SNPs associated with type 2 diabetes and differential DNA methylation in human pancreatic islets. Diabetologia. 56: 1036–46.

34. Dos Santos, C., P. Bougnères, and D. Fradin. 2009. A single-nucleotide polymorphism in a methylatable Foxa2 binding site of the G6PC2 promoter is associated with insulin secretion in vivo and increased promoter activity in vitro. Diabetes. 58: 489–92.

35. Mittelstraß, K. , and M. Waldenberger. 2018. DNA methylation in human lipid metabolism and related diseases. Curr. Opin. Lipidol. 29: 116–124.

36. Holle, R., M. Happich, H. Löwel, H. E. Wichmann, and MONICA/KORA Study Group. 2005. KORA--a research platform for population based health research. Gesundheitswesen. 67 Suppl 1: S19–25.

37. Zeilinger, S., B. Kühnel, N. Klopp, H. Baurecht, A. Kleinschmidt, C. Gieger, S. Weidinger, E. Lattka, J. Adamski, A. Peters, K. Strauch, M. Waldenberger, and T. Illig. 2013. Tobacco smoking leads to extensive genome-wide changes in DNA methylation. PLoS One. 8: e63812.

38. Chen, L. S., and J. D. Storey. 2008. Eigen-R2 for dissecting variation in high-dimensional studies. Bioinformatics. 24: 2260–2.

39. Lehne, B., A. W. Drong, M. Loh, W. Zhang, W. R. Scott, S.-T. Tan, U. Afzal, J. Scott, M.-R. Jarvelin, P. Elliott, M. I. McCarthy, J. S. Kooner, and J. C. Chambers. 2015. A coherent approach for analysis of the Illumina HumanMethylation450 BeadChip improves data quality and performance in epigenome-wide association studies. Genome Biol. 16: 37.

40. Aryee, M. J., A. E. Jaffe, H. Corrada-Bravo, C. Ladd-Acosta, A. P. Feinberg, K. D. Hansen, and R. Irizarry. 2014. Minfi: a flexible and comprehensive Bioconductor package for the analysis of Infinium DNA methylation microarrays. Bioinformatics. 30: 1363–9.

41. Smyth, G. K. 2005. *In* Bioinformatics and Computational Biology Solutions Using R and Bioconductor (Gentleman, R., Carey, V. J., Huber, W., Irizarry, R. A., and Dudoit, S., eds.). pp. 397–420., Springer New York, New York, NY.

42. Touleimat, N., and J. Tost. 2012. Complete pipeline for Infinium(®) Human Methylation 450K BeadChip data processing using subset quantile normalization for accurate DNA methylation estimation. Epigenomics. 4: 325–41.

43. Pfeiffer, L., S. Wahl, L. C. Pilling, E. Reischl, J. K. Sandling, S. Kunze, L. M. Holdt, A. Kretschmer, K. Schramm, J. Adamski, N. Klopp, T. Illig, Å. K. Hedman, M. Roden, D. G. Hernandez, A. B. Singleton, W. E. Thasler, H. Grallert, C. Gieger, C. Herder, D. Teupser, C. Meisinger, T. D. Spector, F. Kronenberg, H. Prokisch, D. Melzer, A. Peters, P. Deloukas, L. Ferrucci, and M. Waldenberger. 2015. DNA Methylation of Lipid-Related Genes Affects Blood Lipid Levels. Circ. Cardiovasc. Genet. 8: 334–342.

44. Kuhn, R. M., D. Karolchik, A. S. Zweig, T. Wang, K. E. Smith, K. R. Rosenbloom, B. Rhead, B. J. Raney, A. Pohl, M. Pheasant, L. R. Meyer, F. Hsu, A. S. Hinrichs, R. A. Harte, B. Giardine, P. A. Fujita, M. Diekhans, T. R. Dreszer, H. Clawson, G. P. Barber, D. Haussler, W. J. Kent, D. Karolchik, R. M. Kuhn, A. S. Hinrichs, A. S. Zweig, P. A. Fujita, M. Diekhans, K. E. Smith, K. R. Rosenbloom, B. J. Raney, A. Pohl, M. Pheasant, L. R. Meyer, K. Learned, F. Hsu, J. Hillman-Jackson, R. A. Harte, B. Giardine, T. R. Dreszer, H. Clawson, G. P. Barber, D. Haussler, and W. J. Kent. 2009. The UCSC Genome Browser Database: update 2009. Nucleic Acids Res. 37: D613–D619.

45. van Oosterhout, C., W. F. Hutchinson, D. P. M. Wills, and P. Shipley. 2004. micro-checker: software for identifying and correcting genotyping errors in microsatellite data. Mol. Ecol. Notes. 4: 535–538.

46. Excoffier, L., G. Laval, and S. Schneider. 2005. Arlequin (version 3.0): an integrated software package for population genetics data analysis. Evol. Bioinform. Online. 1: 47–50.

47. Kronenberg, F., E. M. Lobentanz, P. König, G. Utermann, and H. Dieplinger. 1994. Effect of sample storage on the measurement of lipoprotein[a], apolipoproteins B and A-IV, total and high density lipoprotein cholesterol and triglycerides. J. Lipid Res. 35: 1318–28.

48. Houseman, E. A., W. P. Accomando, D. C. Koestler, B. C. Christensen, C. J. Marsit, H. H. Nelson, J. K. Wiencke, and K. T. Kelsey. 2012. DNA methylation arrays as surrogate measures of cell mixture distribution. BMC Bioinformatics. 13: 86.

49. GTEx Consortium. 2015. Human genomics. The Genotype-Tissue Expression (GTEx) pilot analysis: multitissue gene regulation in humans. Science. 348: 648–60.

50. Matys, V., O. V Kel-Margoulis, E. Fricke, I. Liebich, S. Land, A. Barre-Dirrie, I. Reuter, D. Chekmenev, M. Krull, K. Hornischer, N. Voss, P. Stegmaier, B. Lewicki-Potapov, H. Saxel, A. E. Kel, and E. Wingender. 2006. TRANSFAC and its module TRANSCompel: transcriptional gene regulation in eukaryotes. Nucleic Acids Res. 34: D108–D110.

51. Mele, M., P. G. Ferreira, F. Reverter, D. S. DeLuca, J. Monlong, M. Sammeth, T. R. Young, J. M. Goldmann, D. D. Pervouchine, T. J. Sullivan, R. Johnson, A. V. Segre, S. Djebali, A. Niarchou, T. G. Consortium, F. A. Wright, T. Lappalainen, M. Calvo, G. Getz, E. T. Dermitzakis, K. G. Ardlie, and R. Guigo. 2015. The human transcriptome across tissues and individuals. Science (80-.). 348: 660–665.

52. Tsimikas, S. 2017. A Test in Context: Lipoprotein(a): Diagnosis, Prognosis, Controversies, and Emerging Therapies. J. Am. Coll. Cardiol. 69: 692–711.

53. Coassin, S., G. Erhart, H. Weissensteiner, M. Eca Guimarães de Araújo, C. Lamina, S. Schönherr, L. Forer, M. Haun, J. L. Losso, A. Köttgen, K. Schmidt, G. Utermann, A. Peters, C. Gieger, K. Strauch, A. Finkenstedt, R. Bale, H. Zoller, B. Paulweber, K.-U. Eckardt, A. Hüttenhofer, L. A. Huber, and F. Kronenberg. 2017. A novel but frequent variant in LPA KIV-2 is associated with a pronounced Lp(a) and cardiovascular risk reduction. Eur. Heart J. 38: 1823–1831.

54. Ogorelkova, M., a Gruber, and G. Utermann. 1999. Molecular basis of congenital lp(a) deficiency: a frequent apo(a) “null” mutation in caucasians. Hum. Mol. Genet. 8: 2087–96.

55. Puckey, L. H., R. M. Lawn, and B. L. Knight. 1997. Polymorphisms in the apolipoprotein(a) gene and their relationship to allele size and plasma lipoprotein(a) concentration. Hum. Mol. Genet. 6: 1099–107.

56. Moriarty, P. M., S. A. Varvel, P. L. S. M. Gordts, J. P. McConnell, and S. Tsimikas. 2017. Lipoprotein(a) Mass Levels Increase Significantly According to APOE Genotype: An Analysis of 431 239 Patients. Arterioscler. Thromb. Vasc. Biol. ATVBAHA.116.308704.

57. Taliun, D., D. N. Harris, M. D. Kessler, J. Carlson, Z. A. Szpiech, R. Torres, S. A. G. Taliun, A. Corvelo, S. M. Gogarten, H. M. Kang, A. N. Pitsillides, J. LeFaive, S. Lee, X. Tian, B. L. Browning, S. Das, A.-K. Emde, W. E. Clarke, D. P. Loesch, A. C. Shetty, T. W. Blackwell, Q. Wong, F. Aguet, C. Albert, A. Alonso, K. G. Ardlie, S. Aslibekyan, P. L. Auer, J. Barnard, R. G. Barr, L. C. Becker, R. L. Beer, E. J. Benjamin, L. F. Bielak, J. Blangero, M. Boehnke, D. W. Bowden, J. A. Brody, E. G. Burchard, B. E. Cade, J. F. Casella, B. Chalazan, Y.-D. I. Chen, M. H. Cho, S. H. Choi, M. K. Chung, C. B. Clish, A. Correa, J. E. Curran, B. Custer, et al. 2019. Sequencing of 53,831 diverse genomes from the NHLBI TOPMed Program. bioRxiv. 563866.

58. Ma, B., E. H. Wilker, S. A. G. Willis-Owen, H.-M. Byun, K. C. C. Wong, V. Motta, A. A. Baccarelli, J. Schwartz, W. O. C. M. Cookson, K. Khabbaz, M. A. Mittleman, M. F. Moffatt, and L. Liang. 2014. Predicting DNA methylation level across human tissues. Nucleic Acids Res. 42: 3515–28.

59. Fan, S., and X. Zhang. 2009. CpG island methylation pattern in different human tissues and its correlation with gene expression. Biochem. Biophys. Res. Commun. 383: 421–5.

60. Lanktree, M. B., S. S. Anand, S. Yusuf, and R. a Hegele. 2010. Comprehensive analysis of genomic variation in the LPA locus and its relationship to plasma lipoprotein(a) in South Asians, Chinese, and European Caucasians. Circ. Cardiovasc. Genet. 3: 39–46.

61. Schmidt, K., A. Noureen, F. Kronenberg, and G. Utermann. 2016. Structure, function, and genetics of lipoprotein (a). J. Lipid Res. 57: 1339–1359.

