## Supplemental Tables and Figures for "A genome-wide analysis of DNA methylation identifies a novel association signal for Lp(a) concentrations in the *LPA* promoter"

**Running title:** Genome-wide methylation study on Lp(a)

Stefan Coassin[1], Natascha Hermann-Kleiter[2], Margot Haun[1], Simone Wahl[3,5], Rory Wilson[3,5], Bernhard Paulweber[4], Sonja Kunze[3,5], Thomas Meitinger[6,7,8], Konstantin Strauch[9,10], Annette Peters[5,6], Melanie Waldenberger[3,5,6], Florian Kronenberg[1,] Claudia Lamina[1]

[1]Institute of Genetic Epidemiology, Department of Genetics and Pharmacology, Medical University of Innsbruck, Austria
[2]Institute of Cell Genetics, Department of Genetics and Pharmacology, Medical University of Innsbruck, Austria
[3]Research Unit of Molecular Epidemiology, Helmholtz Zentrum München – German Research Center for Environmental Health, Neuherberg, Germany
[4]First Department of Internal Medicine, Paracelsus Private Medical University, Salzburg, Austria
[5]Institute of Epidemiology II, Helmholtz Zentrum München – German Research Center for Environmental Health, Neuherberg, Germany
[6]German Center for Cardiovascular Research (DZHK), Partner Site Munich Heart Alliance, Munich, Germany
[7]Institute of Human Genetics, Technische Universität München, Munich, Germany
[8]Institute of Human Genetics, Helmholtz Zentrum München, German Research Center for Environmental Health, Neuherberg, Germany
[9]Institute of Genetic Epidemiology, Helmholtz Zentrum München, German Research Center for Environmental Health, Neuherberg, Germany.
[10]Institute of Medical Informatics, Biometry, and Epidemiology, Ludwig-Maximilians-Universität, Munich, Germany.

**Address of correspondence:**
Claudia Lamina
Institute of Genetic Epidemiology
Department of Genetics and Pharmacology
Medical University of Innsbruck
Schöpfstr. 41, A-6020 Innsbruck, AUSTRIA
Phone: (+43) 512 9003 70565


### Supplementary Tables

**Supplementary Table 1:** CpG sites in the *LPA* gene locus assayed by the Illumina Infinium HumanMethylation450 BeadChip Array with position, location and the p-value of the first stage epigenome-wide analysis on log(Lp(a)). The location is given relative to the UCSC Genome browser hg38 annotation with a leading non-coding exon 1. See the introduction section of the main manuscript for an explanation of the issues with the *LPA* reference transcript annotation.

| cg-ID | Chr | Position | location | p-value |
|---|---|---|---|---|
| cg10234069 | 6 | 160952713 | Exon 40 | 0.1193 |
| cg14663533 | 6 | 160955178 | Intron 38-39 | 0.9046 |
| **cg17028067** | **6** | **161086715** | **Intron 1-2** | **6.04e-11** |
| cg10836120 | 6 | 161087339 | Exon 1 | 0.1954 |
| cg11975608 | 6 | 161087439 | 5´upstream | 0.8437 |
| cg17189167 | 6 | 161087506 | 5´upstream | 0.4256 |
| cg16960593 | 6 | 161087573 | 5´upstream | 0.4193 |
| cg27656787 | 6 | 161088246 | 5´upstream | 0.3816 |
| cg07177174 | 6 | 161088755 | 5´upstream | 0.0104 |

**Supplementary Table 1:** Primer sequences. For Bisulfite Sanger sequencing primer the lower case letters denote tails that have been added by the design software to increase the annealing temperature of the primer. The PCR protocols are given in the Methods section of the manuscript. See Supplementary Figure 2 for an overview of the target sequence.

| **Primer-ID** | **Sequence (5' – 3')** | **use** | **annealing temperature [°C]** |
|---|---|---|---|
| LPA_as_785_fw | aggaagagagTAGGAGGTGGAAGTTGTAGTGAGTT | bisulfite sequencing amplicon 1 and sequencing primer | 66 |
| LPA_as_323_rv | cagtaatacgactcactatagggagaaggctAAACCACTCACCCTCCTAAAATATC | bisulfite sequencing amplicon 1 | 66 |
| LPA_sense_1211_fw | aggaagagagTGGGATGATTGGTATGTGTTTTTAT | bisulfite sequencing amplicon 2 and sequencing primer | 63 |
| LPA_sense_1694_rv | cagtaatacgactcactatagggagaaggctCCAAACCCTCTACCAAATACTACAC | bisulfite sequencing amplicon 2 | 63 |
| LPA_fw_rs76735376 | TACAGGACAGAGACTAACT | amplicon for validation of rs76735376 by sequencing | 60 |
| LPA_rw_rs76735376 | GCATAGTATCAATCTTTCCG | amplicon for validation of rs76735376 by sequencing | 60 |
| LPA_sense_1377_fw | TTGGAAGGATTGATATTTTATAATATAATTTAT | sequencing primer for bisulfite DNA for amplicon 2 | 55 |
| LPA_as_408_rv | CAAAAACTAACTACACACAATATCTAAAACT | sequencing primer for bisulfite DNA for amplicon 1 | 55 |

**Supplementary Table 3:** Study characteristics (Mean ±sd [25%,50%,75% Percentile] for quantitative variables, n(%) for gender

| Study | KORA F4<br>n=2986*<br>n=1724† | KORA F3<br>n=3080*<br>n=484† | SAPHIR<br>n=1446* |
|---|---|---|---|
| Age, yrs | 56.1±13.3 [44,56,67]<br>61.0±8.9 [54,61,68] | 57.4±12.9 [46.8,57,67]<br>53.2±9.6 [46,54,61] | 51.0±6.0 [46,52,55] |
| Gender (female) | 1545 (51.7%)<br>881 (51.1%) | 1583 (51.4%)<br>232 (47.9%) | 466 (32.2%) |
| Lp(a), mg/dL | 21.7±24.6 [5.2,11.7,30.3]<br>22.4±25.1 [5.5,12.1,31.3] | 22.1±26.2 [4.9,11.2,28.6]<br>20.1±23.1 [5.2,10.2,25.2] | 23.9±27.4<br>[5.3,11.6,35.6] |
| Apo(a) isoforms, KIV repeats‡ | 26.8±6.0 [23,26,31]<br>26.8±5.9 [23,26,31] | 27.2±6.0 [23,27,31]<br>27.1±5.8 [23,27,31] | 26.9±6.2 [22,26,31] |

*First line in each cell: dataset available for SNP association analysis (with non-missing genotype, age, sex and Lp(a));
†Second line in each cell: subset available for methylation analysis (with no missing values in primary linear regression model);
‡ in heterozygous individuals the smaller of both isoforms per person is used

**Supplementary Table 4:** Genotype frequencies of the de novo genotyped SNP rs76735376 in three cohorts.

| Study | Number of genotypes and frequencies<br>CC | <br>CT | <br>TT | Minor allele frequency |
|---|---|---|---|---|
| KORA F4 | 2912 (0.975) | 74 (0.025) | 0 (0) | 0.012 |
| KORA F3 | 3016 (0.979) | 63 (0.020) | 1 (0.0003) | 0.011 |
| SAPHIR | 1419 (0.981) | 27 (0.019) | 0 (0) | 0.009 |

**Supplementary Table 5:** Allele frequencies of the PNR in SAPHIR and KORA F4, separated for genotypes of rs76735376.

| | SAPHIR | | KORA F4 | |
|---|---|---|---|---|
| PNR allele [n repeats] | PNR allele Carriers, n (%) in CC-carriers | PNR allele Carriers, n (%) in CT-carriers | PNR allele Carriers, n (%) in CC-carriers | PNR allele Carriers, n (%) in CT-carriers |
| 4 | 1 (0.07) | 0 (0) | 0 (0) | 0 (0) |
| 5 | 0 (0.00) | 0 (0) | 5 (0.17) | 0 (0) |
| 6 | 7 (0.50) | 0 (0) | 4 (0.14) | 0 (0) |
| 7 | 14 (1.00) | 0 (0) | 19 (0.66) | 0 (0) |
| 8 | 1250 (89.16) | 27 (100) | 2570 (89.17) | 74 (100) |
| 9 | 103 (7.35) | 0 (0) | 202 (7.01) | 0 (0) |
| 10 | 27 (1.92) | 0 (0) | 82 (2.84) | 0 (0) |

#### Supplementary Figures

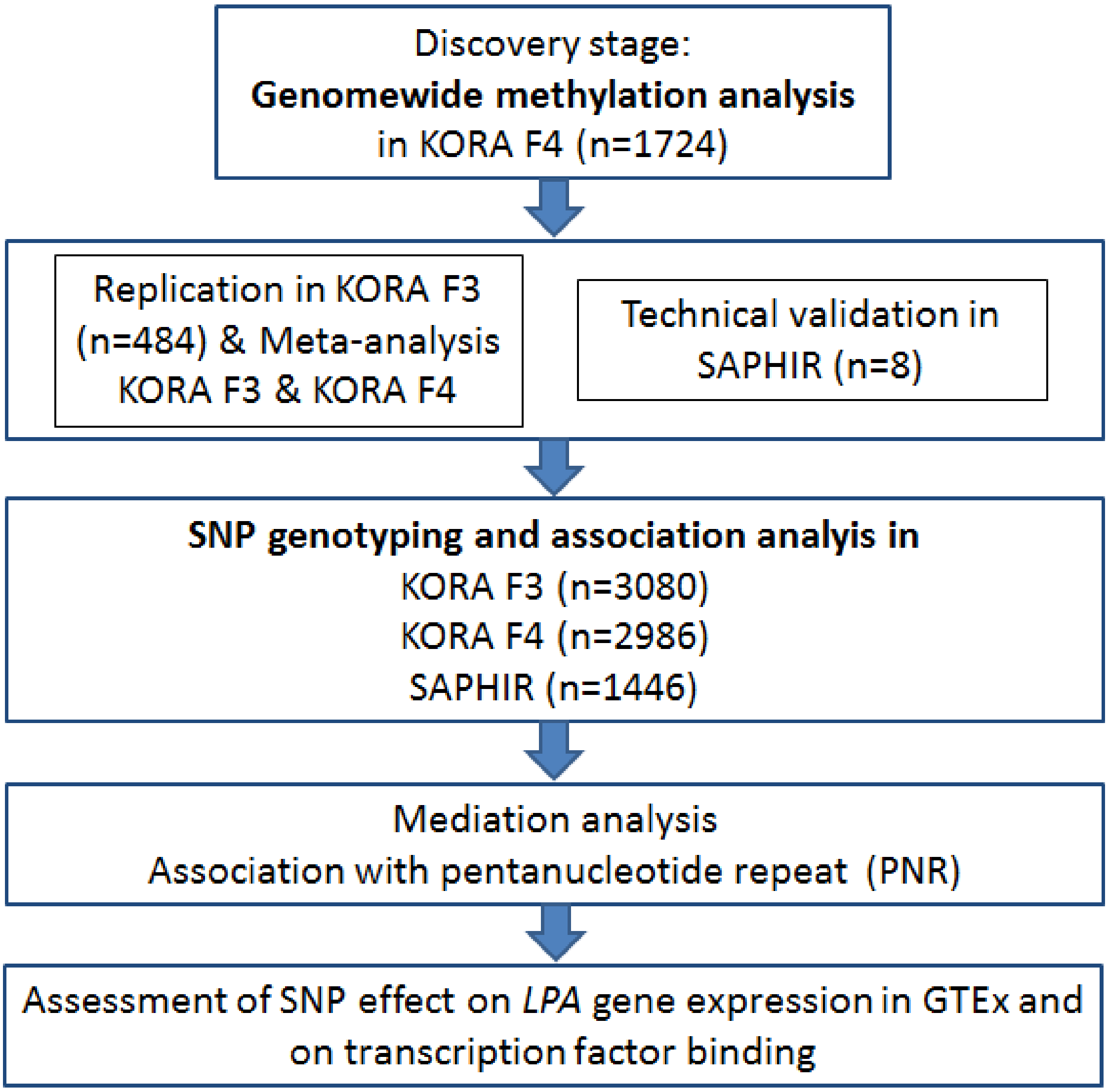


**Supplementary Figure 1:** Flow chart of the study design

ctatgagtttctgaatggaagagtcaaatccactgagctctgtgctgggcatttggctggaggcattg

acagttgcaaggtaagaaaagatcaagagaccaaagttagtcttgtgctctcctgtctcagtctcagt

cccttagacttgagtcccaaagtagcgaattcaagtaggatttaatcaatggaagaccccagtctaag

tgttgctcagaaactccctagatctgtcccaaatgtatattcagatcatccaaggggacttcttgggg

cttgagttccagatcagcagcaagggagccataagtgccataactacctcAGACCACTCACCCTCCTG

GGGTGTCCCGGTGGCCAGGGACTAAAGTGGTGATTTTTCTGGTAGGGAAGGAGGTAGAGGGTACAGGA

CAGAGACTAACTGCACACAATATCTGAGACTGGAGCTCAGATATTGCTGATGATCAGAGTTGGCGTGT

CTCCCCAATTGATTTACAACTGGGGCTTGGATACTGTTTTAAACGGGAGGAGCCTCCTAACCATCTTG

ACACAACCACTGACGTGACTACACTAGAATAGACTCTTTCCACTTAATTCTACCACTCTTGCTTTACT

TCATGAGAACGAAAATGTAAGATTGCACCATGAATTCATTTGCGGAAAGATTGATACTATGC**TTTTAT**

**TTTATTTTATTTTATTTTATTTTATTTTATTTTATTTTAT**TGAGACTCTCACCCCGGTTGAAGTGCAC

TGACGTGATTTTGGCTCACTGCAACTTCCACCTCCTgggttcaagtgaatactccagcctccctagta

gctgggattacaggtgcccaccaccacgcctggctaattttgtatttttagtagagatggggtttca

ccacattggcctggctggtctcaaactcctgaccttgtgatccacctgtcttggcctcccaaagtgct

gggattacagagttgagccaccgcactcgaccctatgtttattttaaaatatttatttatttatt

taagccacaactactagaataggaaggattgatattttattaattttatttggtatttattattttt

tttctttcctgagacattcttgctctgtcacccaggctggagtgcagtggcacattcttggctcactg

caacctccatctcctgtgttcaagcaattctagtgcctcagcctacttagtagcTGGGATGACTGGCA

TGTGCCTCCACACCCAGCTAATTTTTGTATTTTTTGTAGAGACAGGGTTTTGGCATGTTGCCCAGGCT

TGTCTCAAACTCCTGGCCTCAGGTGATCCATCTGCCGTGGCCTCCCAAAATGCTGGGATTATAGGCAT

GAGCCACCACCCCCTCCTGGAAGGATTGATATCTTATAACATAATTTATAATTACAGAAAACATGTGA

GTTCACTAGGAATAAATAAATTTTGAAGATAATAAAAGATTTTCACTTATGTTGTCATTTCGGCACAG

TTTGGTATAGGATGTGGAGATGTTAACATTTATACCTAGCTTGCTCGTAAACTAAGACCTGAAAGGGT

TGTGTCTATCAGCTGCACCCCTGGGTAGCGACACAACCTCGGGAAGGCCTCAGCCCCCTCCTCGTACA

GCACTGCCTGTTGGAAAGCTTGAGGGAGGCTATGGATGTGCAGCACTTGGCAGAGGGTCTGGtcatgg

aagttaccagcaaatatgagctacttttatgattttattttatccaaaagaaagagaatgaaagaaga

ggggaggaaacaagactaatcaggaaagatgaaggtctaggggtgagggaaggagtaaggagacataa

aggcaatgtggagcagctgagggggaaatggctttcaccacttcccagcatctattgacattgcact

ctcaaatattttataagactctatattcaaggtaatgtttgaaccctgctgagccagtggcatgggtc

tctgagagaatcattaacttaatttgactatctggtttgtgggtgcgtttactctcatgtaagtcaa

**Supplementary Figure 2**: Overview of the region around rs76735376 with location of the LPA pentanucleotide repeat, the location of the amplicons, the location of the CpG affected by rs76735376 and neighboring CpG sites.

Upper case: sequence regions, which could be covered with amplicons for bisulfite sequencing. Lower case: regions which could not be covered by amplicons for bisulfite sequencing.

Blue background (light and dark): primer binding sites for bisulfite sequencing. Note that the sequence given here is the unconverted sequence. The primer given in Supplementary Table 2 bind on the converted sequence and the complement strand of the strand shown here. Yellow background: primer binding sites of the sequences used for SNP validation by sequencing. Bold character: *LPA* pentanucleotide repeat. Green: CpG affected by rs76735376 (enlarged cytosine base). Position of rs76735376 is underlined. Pink: CpG in the region, which were amenable to sequencing. Purple: CpG in the region, which were not amenable to sequencing (not represented in any amplicon).

Sequence is the bisulfite-converted sequence given 5’ to 3’ on the minus strand of human genome GRCh37/hg19 (i.e. the same direction as LPA transcription direction).

```
Oligo2 LPA_EMSA2-C-f      5'-CATGGTGCAATCTTACATTTTCGTTCTCAT-3'
Oligo4 LPA_EMSA2-T-f      5'-CATGGTGCAATCTTACATTTTCATTCTCAT-3'
Oligo6 LPA_EMSA2-5mC-f    5'-CATGGTGCAATCTTACATTTTCGTTCTCAT -3'

Oligo2 LPA_EMSA2-C-r      3'-GTACCACGTTAGAATGTAAAAG CAAGAGTA-5'
Oligo4 LPA_EMSA2-T-r      3'-GTACCACGTTAGAATGTAAAAG TAAGAGTA-5'
Oligo6 LPA_EMSA2-5mC-r    3'-GTACCACGTTAGAATGTAAAAGmCAAGAGTA-5'
```

**Supplementary Figure 3:** Oligos used for EMSA experiments. The bold case base gives the location of rs76735376. Green background: POU2F1/POU5F1 binding site, Green font: mutated POU2F1/POU5F1 binding site, Blue CEBPB binding site. The reverse primers (-r) are given in reverse orientation, as they are annealed to the forward oligos (-f).

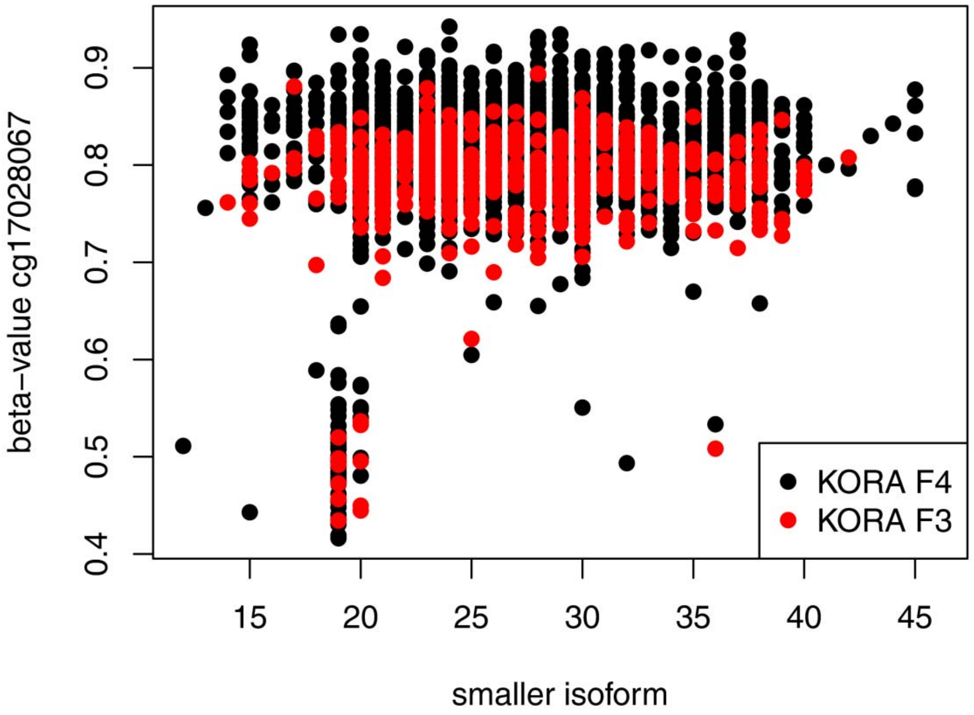


**Supplementary Figure 4:** Scatterplot showing the lower of both apo(a) isoforms (x-axis) per person in KORA F3 (in red) and KORA F4 (in black) versus the beta-value of cg17028067 (y-axis)

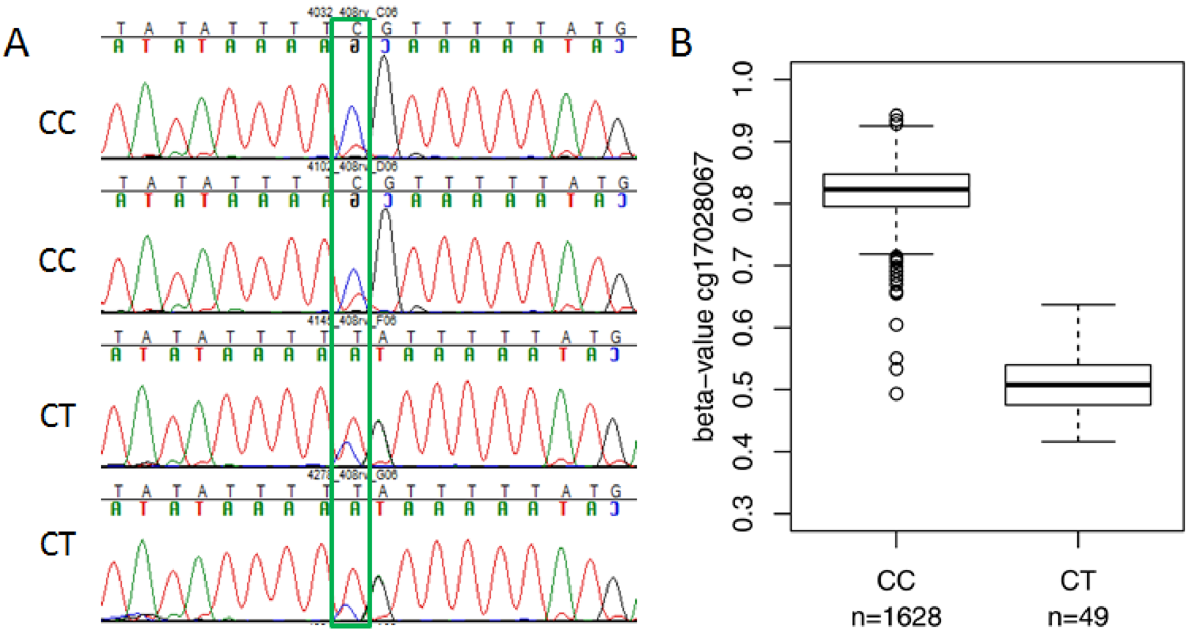


**Supplementary Figure 5: Panel A:** Representative results of bisulfite sequencing in two homozygotes for the major allele and two heterozygotes in the SAPHIR study. The blue peak represents the unconverted C-allele, indicating that the major part is methylated in CC carriers, whereas only a minor part is methylated in CT carriers. **Panel B:** Boxplots of the methylation level (expressed as beta-value) of cg17028067, stratified for genotypes in the KORAF4 study (panel B).

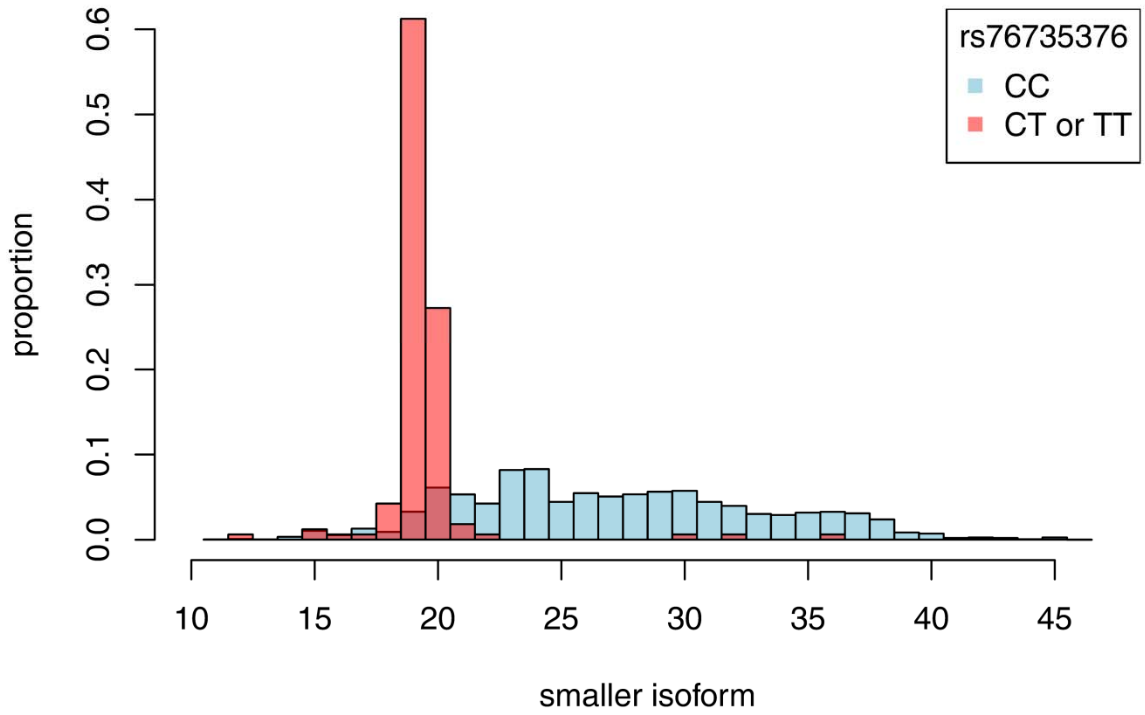


**Supplementary Figure 6:** Distribution of the smaller of both isoforms, stratified for carriers (CT or TT) and non-carriers (CC) of the SNP rs76735376.

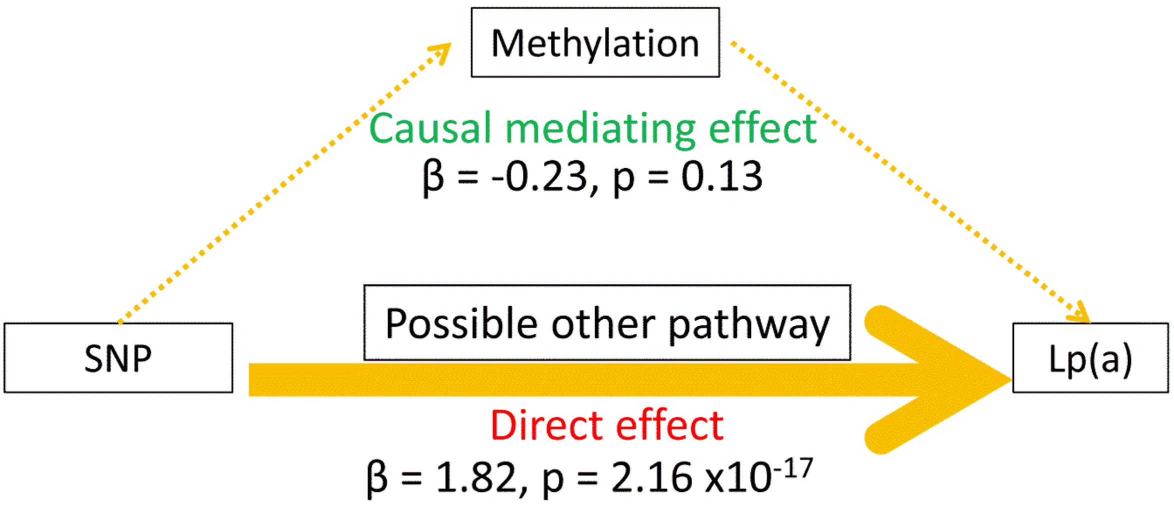


**Supplementary Figure 7:** Possible paths and results of mediation analysis.

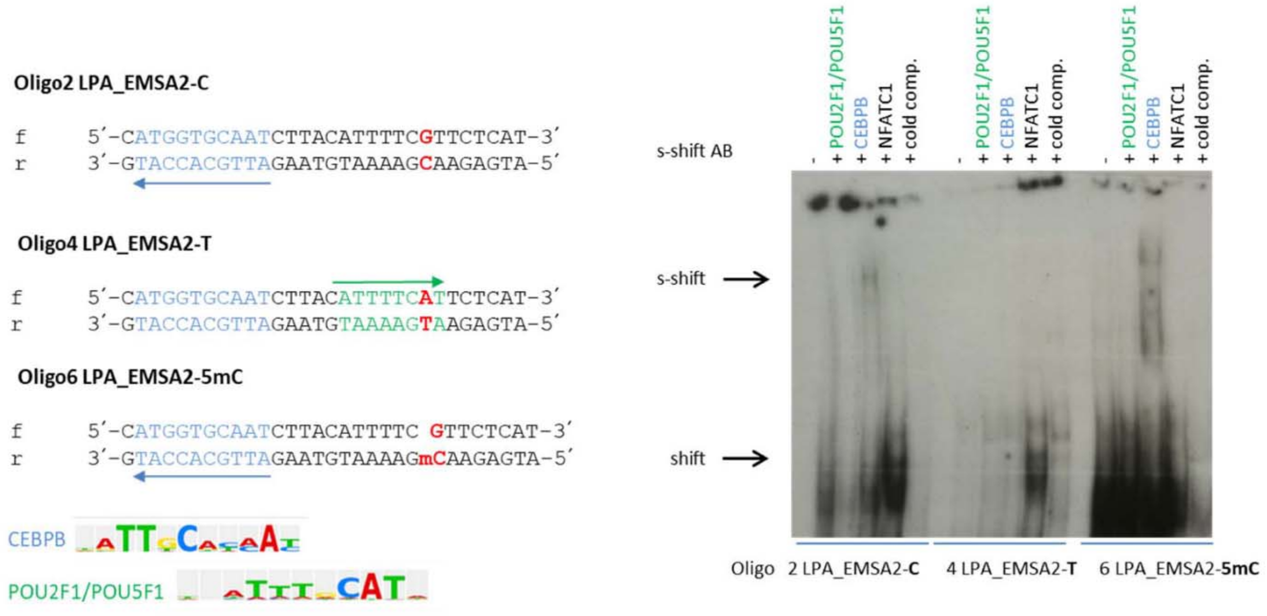


**Supplementary Figure 8: Electrophoretic mobility shift assay for rs76735376**

rs76735376 modifies binding of the transcription factor CEBPB. Left panel: Diagram of the double-stranded oligos used for EMSA analysis containing either cytosine (red, Oligo2 LPA_EMSA2-C), thymine (Oligo4 LPA_EMSA2-T) or methyl-cytosine (6 LPA_EMSA2-5mC) oligo. The transcription factor binding sites for CEBPB (blue) and POU2F1/POU5F1 (green) as predicted by Transfac analysis, 52 of the rs76735376 probes are shown. Position weight matrix of the TF is shown. The arrow indicates the orientation of the sequence matrix in the oligo. Right panel: EMSA analysis of human liver nuclear extract hybridized to Oligo2 LPA_EMSA2-C, Oligo4 LPA_EMSA2-T or Oligo6 LPA_EMSA2-5mC. Binding specificity to specific transcription factors was performed by using super-shift antibodies (s-shift AB) for the POU factors or CEBPB. An NFATC1 antibody and excess of cold unlabeled oligonucleotide (cold comp.) were used as negative control. One representative experiment out of three is shown.

Note that the free probe has already left the gel because a long run time in order was required to distinguish the super-shift bands.
